# A new framework for *Subti*Wiki, the database for the model organism *Bacillus subtilis*

**DOI:** 10.1101/2024.09.10.612211

**Authors:** Christoph Elfmann, Vincenz Dumann, Tim van den Berg, Jörg Stülke

**Affiliations:** Department of General Microbiology, Institute of Microbiology and Genetics, Georg-August University Göttingen, Grisebachstr. 8, D-37077 Göttingen, Germany

**Keywords:** *Subti*Wiki, genome annotation, protein-protein interaction, gene order conservation

## Abstract

*Bacillus subtilis* is a Gram-positive model bacterium and one of the most-studied and best understood organisms. The complex information resulting from its investigation is compiled in the database *Subti*Wiki (https://subtiwiki.uni-goettingen.de/v5) in an integrated and intuitive manner. To enhance the utility of *Subti*Wiki, we have added novel features such as a viewer to interrogate conserved genomic organization, a widget that shows mutant fitness data for all non-essential genes, and a widget showing protein structures, structure predictions and complex structures. Moreover, we have integrated metabolites as new entities. The new framework also includes a documented API, enabling programmatic access to data for computational tasks. Here we present the recent developments of *Subti*Wiki and the current state of the data for this organism.

**Key points:**

- *Subti*Wiki is the most popular database for the model bacterium *Bacillus subtilis*.
- To facilitate the development of new research hypotheses, *Subti*Wiki has been enhanced by graphical information on conserved genomic organization, mutant fitness data, and a new protein structure viewer.
- Metabolites have been integrated as entities on their own with dedicated pages, interactive information on reactions and metabolite functions as well as their interactions.

## Introduction

After decades of intensive research, the functions for a large number of proteins have not yet been identified. Instead, research is focussing on proteins with known functions, and more and more studies describing new details of identified proteins are published each year. In contrast, only little research activity is devoted to the analysis of unknown proteins – collectively referred to as the unknowme. As a result, roughly one third of all proteins in well-studied organisms has no function assigned to them. Even in model organisms such as the bacteria *Escherichia coli* and *Bacillus subtilis*, about 25% of all proteins lack a known function (1, 2). On one hand, this may reflect the intensive investigation of cellular standard functions such as cell division or metabolism, the components of which have mostly been identified. Thus, unknown proteins are often required under highly specific and poorly studied conditions and have therefore escaped the discovery of their functions (3). On the other hand, some of the unknown proteins are strongly expressed under a variety of different conditions and are conserved in many organisms. It is obvious that the function of proteins of the latter class should be studied with highest priority (4). A major problem in the investigation of unknown proteins is the lack of hypotheses that could be verified. To develop such hypotheses it is essential that all available information is brought together and made accessible in an intuitive manner. Biological databases are very powerful in the integration of information. As most research is performed on model organisms, the development and continuous improvement of the corresponding model organism databases is highly important. This is true not only for the community that studies the particular organism, but for a broader audience of researchers interested in more or less related species.

We have developed *Subti*Wiki, the database for the Gram-positive model bacterium *Bacillus subtilis* (5, 6). Next to *E. coli* and yeast, *B. subtilis* is the most intensively studied and best understood microorganism (7). Moreover, the research on *B. subtilis* is highly relevant to improve our understanding of important pathogens, among them *Enterococcus faecium* and *Staphylococcus aureus*, that have been listed among the bacterial ESKAPE priority pathogens by the World Health Organization (WHO) due to their widespread resistances against antibiotics (8). Moreover, *B. subtilis* and many of its close relatives are important work horses in a large variety of biotechnological applications, from the production of secreted proteins to the fermentation of vitamins and antibiotics (7).

*Subti*Wiki integrates a wide array of data in an intuitve and interactive manner. This includes data from biochemical or genetic studies on individual genes or proteins as well as results from system level studies. With all this integrated information, *Subti*Wiki has become the most popular database in the field. To further improve the value of the database, we have completely rebuilt the software foundation of *Subti*Wiki. By separating the data layer from the presentation layer, information can not only be accessed easily by users of the web interface, but is also retrievable for bioinformatic purposes. Moreover, we have integrated completely new information such as the conserved genomic organization, the relative fitness of mutants, and dedicated pages for metabolites. Finally, we have integrated an interactive viewer for protein structures and structure predictions of individual proteins and protein complexes. We are confident that the new edition of *Subti*Wiki will facilitate the daily work of a large research community.

### *Subti*Wiki entities: gene/ protein and metabolite pages

So far, *Subti*Wiki has focused on the annotation of genes and proteins. In the new version, we have added dedicated pages for the individual metabolites of *B. subtilis*. The user can search for any page using the search box, which can be found on the start page or on each of the individual pages. The search system has been updated and will show the links to the direct (if hit any), but also to gene and metabolite pages that cover the search item in their name.

For the gene pages, we have kept the general design and page layout (Fig. 1). The main information is promptly presented in the form of a table, such as data on locus name, and – in the case of protein-encoding loci – isoelectric point, molecular weight and function as well as relevant database links. The pages provide manually curated interactive information on specific data such as protein structure, protein-protein interactions, regulatory elements and functional annotation. Each gene/ protein is assigned to one or more functional categories and to regulons. Moreover, the pages contain information on biological materials available in the community, laboratories that study the gene or protein, and relevant publications.

**Figure 1.**
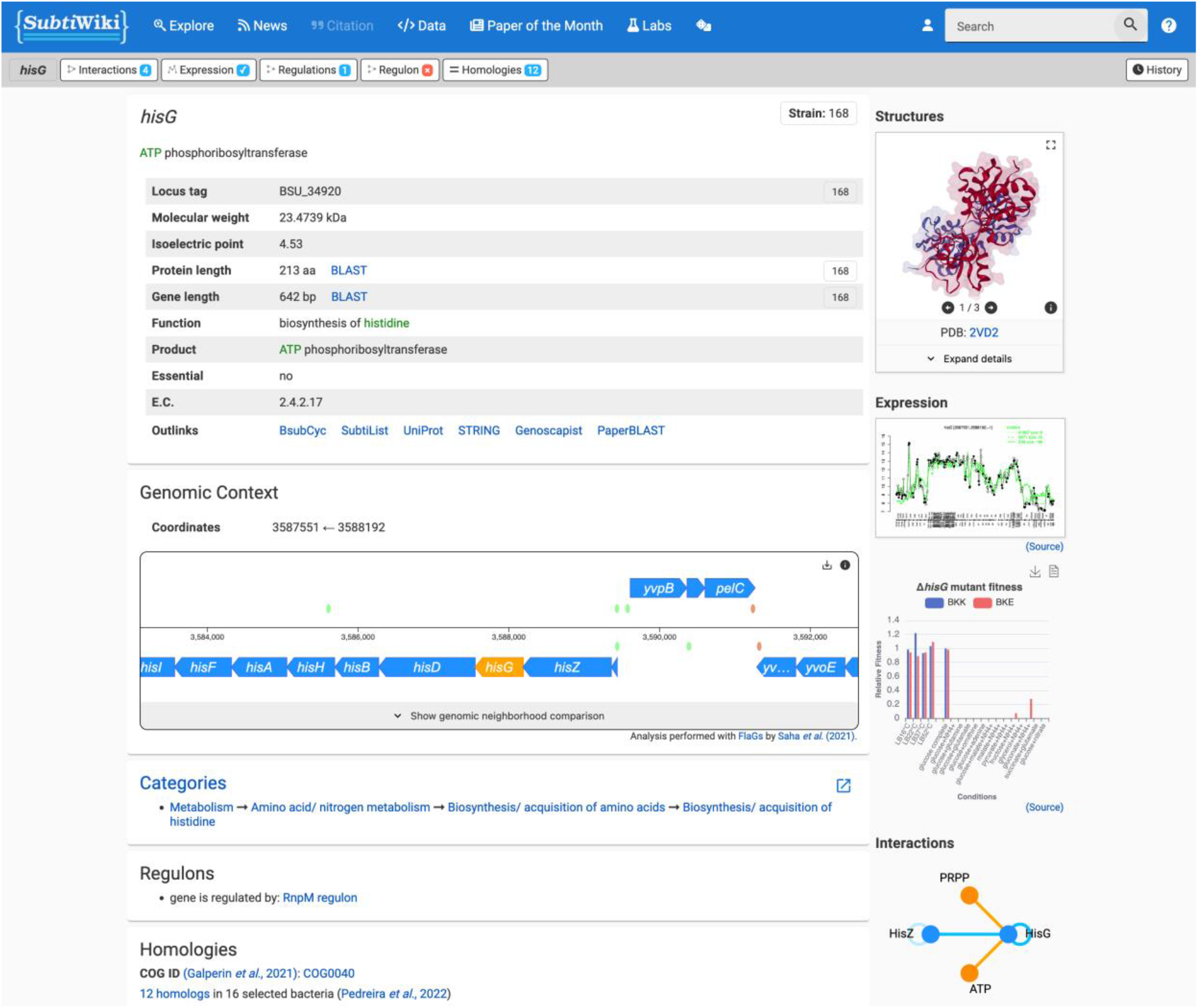
The *hisG* gene page. The page layout has been slightly revised, with a new navigation bar at the top of the page giving access to various related tools and pages. The sidebar now features a 3D visualization of the structure, and a graph showcasing the fitness of the deletion mutant under different conditions. For full access to the page, see https://subtiwiki.uni-goettingen.de/v5/gene/hisG.

A new element on the sidebar of the gene pages is an interactive mutant fitness viewer (9) which displays the relative fitness of mutants for all non-essential genes in complex and selected minimal media using published data (10). Moreover, the interaction widget in the side bar has been extended to interactions with metabolites and other low-molecular-weight molecules.

For the metabolite pages, we have developed an own layout that is similar to that of the gene pages (Fig. 2). These pages feature a table with some key information, lists of reactions the molecule is involved in, its transport (if any), and its role as an effector molecule. As with the gene pages, all information is interactive. Information was retrieved from *Subti*Wiki gene pages and from UniProt database (11). Two widgets in the side bar display the structure of the molecule and its interactions. To distinguish proteins from metabolites, we have introduced a color code with blue for proteins and orange for metabolites for these interaction widgets on both the gene/ protein and metabolite pages.

**Figure 2.**
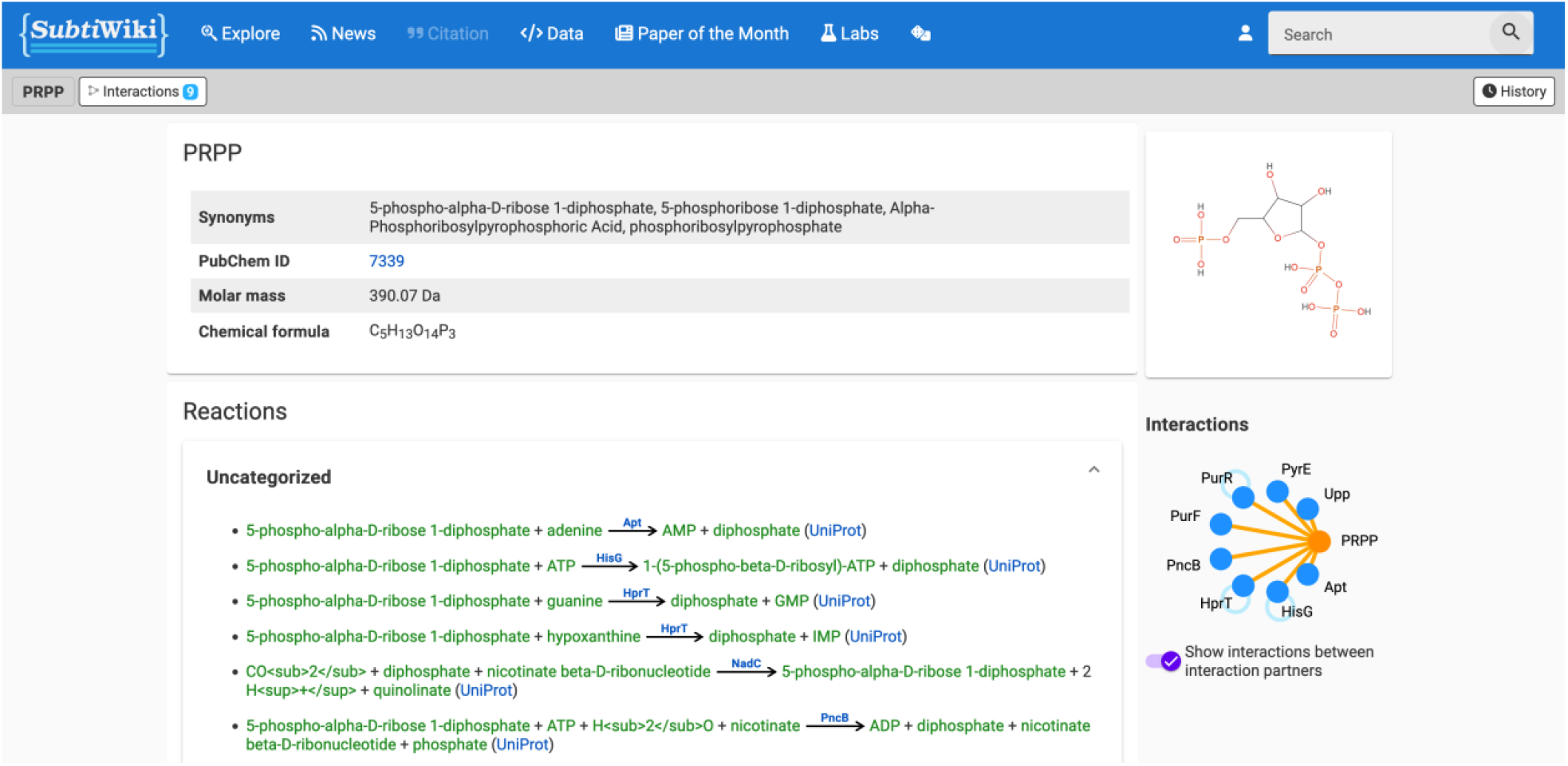
The PRPP metabolite page. Dedicated pages have been created for metabolites, with a layout based on that of gene pages. Basic information is compiled in a table, and additional sections describe the role of the metabolite, listing reactions, transport processes, cofactor interactions, etc. The sidebar shows the molecular structure and a graph of interactions with proteins. For full access to the page, see https://subtiwiki.uni-goettingen.de/v5/metabolite/PRPP.

A novel feature on all pages is a top bar that provides a direct access to interactions, gene expression, regulation, regulons, and homologies.

### Visualization of protein and complex structures and structure predictions

Since the release of the last version of *Subti*Wiki, the application of artificial intelligence using algorithms such as AlphaFold (12) to predict protein structures has revolutionized biology. Accordingly, we have redone the structure widget in the *Subti*Wiki side bar (see Fig. 3), which displays 3D visualizations of molecules using NGL viewer (13). We have now included at least one structure model for each *B. subtilis* protein from the AlphaFold database (14). Moreover, the structure viewer has been greatly enhanced by providing a direct visualization of multiple structure models of a protein and its complexes (if available) in a carrousel-like manner. The structure widget now features a fullscreen mode, and the structures can be displayed in multiple styles including surface charge and hydrophobicity.

**Figure 3.**
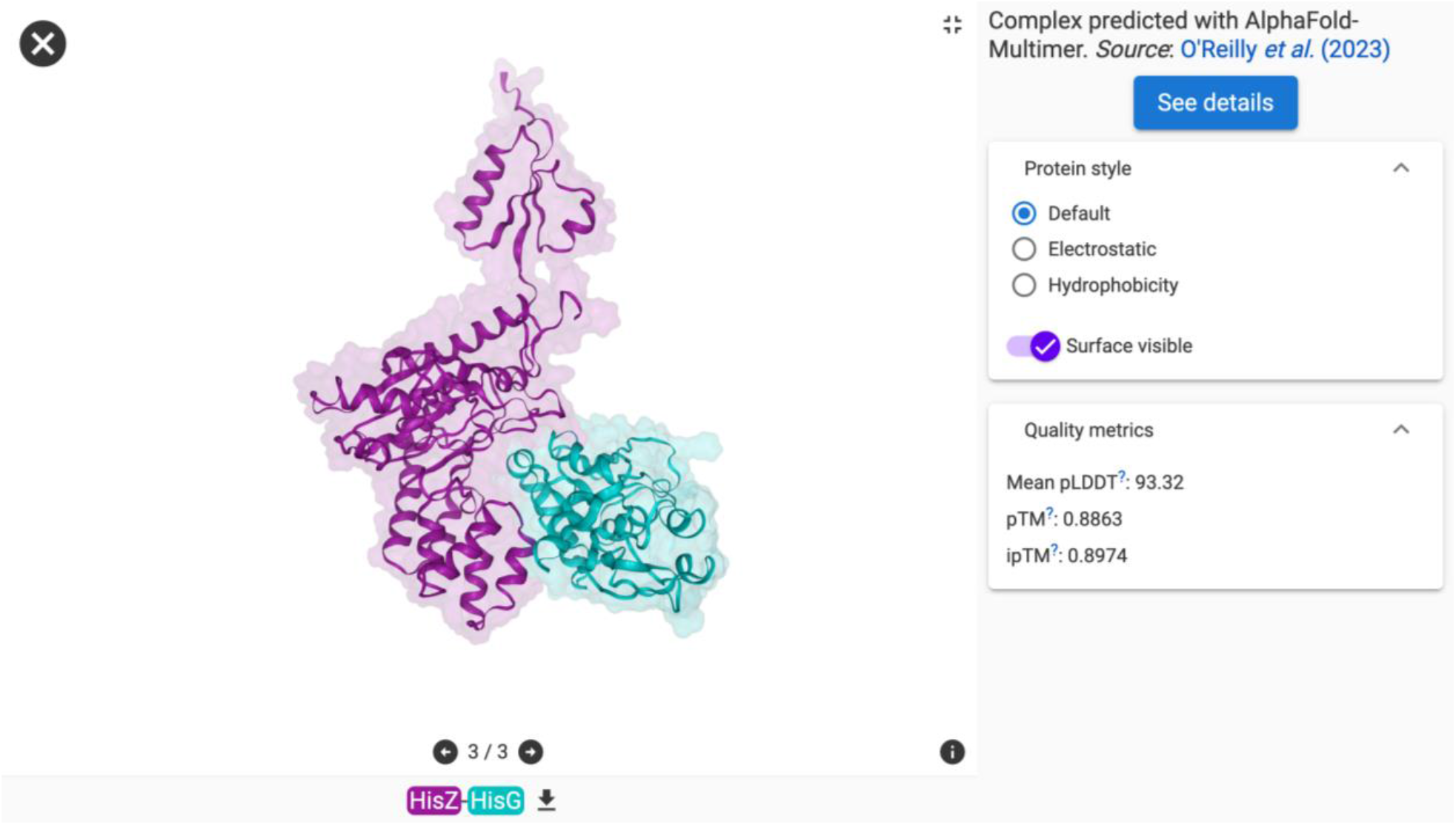
The structure viewer showing the HisG-HisZ complex in fullscreen mode. Interactive 3D visualizations of related structures and structure predictions are now available for each protein.

As many proteins act in complexes, the identification of protein-protein interactions as well as the determination and/or prediction of the complex structures has gained a considerable momentum in recent years. Tools such as AlphaFold-Multimer were developed to facilitate the prediction of complex structures using artificial intelligence (15, 16). Using *in vivo* crosslinking and co-fractionation, a large number of novel protein-protein interactions in *B. subtilis* has recently been identified (17). All these interactions are displayed in the interaction widget in the sidebar as well as in the interaction browser. Moreover, whenever possible, structures of complexes for which no experimental structure was available were predicted using AlphaFold-Multimer (17). All these predictions are also displayed in the structure viewer widget. Importantly, if available, cross-linking information is integrated in these models and it is shown whether a crosslink is in agreement or not with the predicted model. To make this information more accessible for a large community, it is possible to download both the structure data and cross-linking information. Moreover, we have developed the PAE Viewer, a unique web server that helps to evaluate and visualize the quality of complex predictions by taking into account not only the structure prediction, but also the cross-links (18).

Users can adjust the appearance of the structure, investigate the confidence of structure predictions and download corresponding data.

As the established tools for AI-mediated prediction of complex structures are not always reliable, we have integrated experimental cross-links into structure prediction in AlphaLink (19). Even with only one single cross-link, the confidence of the predicted models can be much higher as for the models predicted using AlphaFold-Multimer. As shown for the RpoA-RpoC interaction of RNA polymerase subunits, AlphaFold-Multimer fails to provide a meaningful prediction, whereas AlphaLink predicts a highly confident model that is very similar to the crystal structure. These new predictions will successively be integrated into *Subti*Wiki. In this way, *Subti*Wiki not only serves its own research community but also accompanies the further development of the most fundamental developments of our time that merge structural biochemistry with artificial intelligence.

### Conserved gene neighbourhood

Proteins involved in a common function are often encoded in the same operons in bacteria. Thus, conserved gene neighbourhoods may indicate that two genes are relevant for the same biological process. Furthermore, conservation of gene order can be regarded as a fingerprint of proteins that physically interact with each other (20). Thus, the analysis of gene synteny is very popular to get indications on the function and potential interaction of the encoded proteins (21). We have therefore decided to integrate a novel interactive feature, the genomic neighborhood viewer, as a part of the genomic context viewer (see Fig. 4). To facilitate the analysis for busy experimental scientists who are not used to select among a multitude of options, we have selected a few organisms, more with close and fewer with distant relationship to *B. subtilis*. Using a homology analysis of *B. subtilis* genes (6) as a basis for the FlaGs algorithm (22), we identified clusters of conserved gene neighborhoods. Centered around the initially selected gene, the gene neighborhoods are shown, with conserved gene clusters indicated by a color code. Hovering over any gene in the display will show short functional information from NCBI Protein Database (23). Moreover, a click on the gene will open the NCBI Reference Sequence page for the corresponding protein. Clicking on any *B. subtilis* gene will recenter the display around the chosen gene.

**Figure 4.**
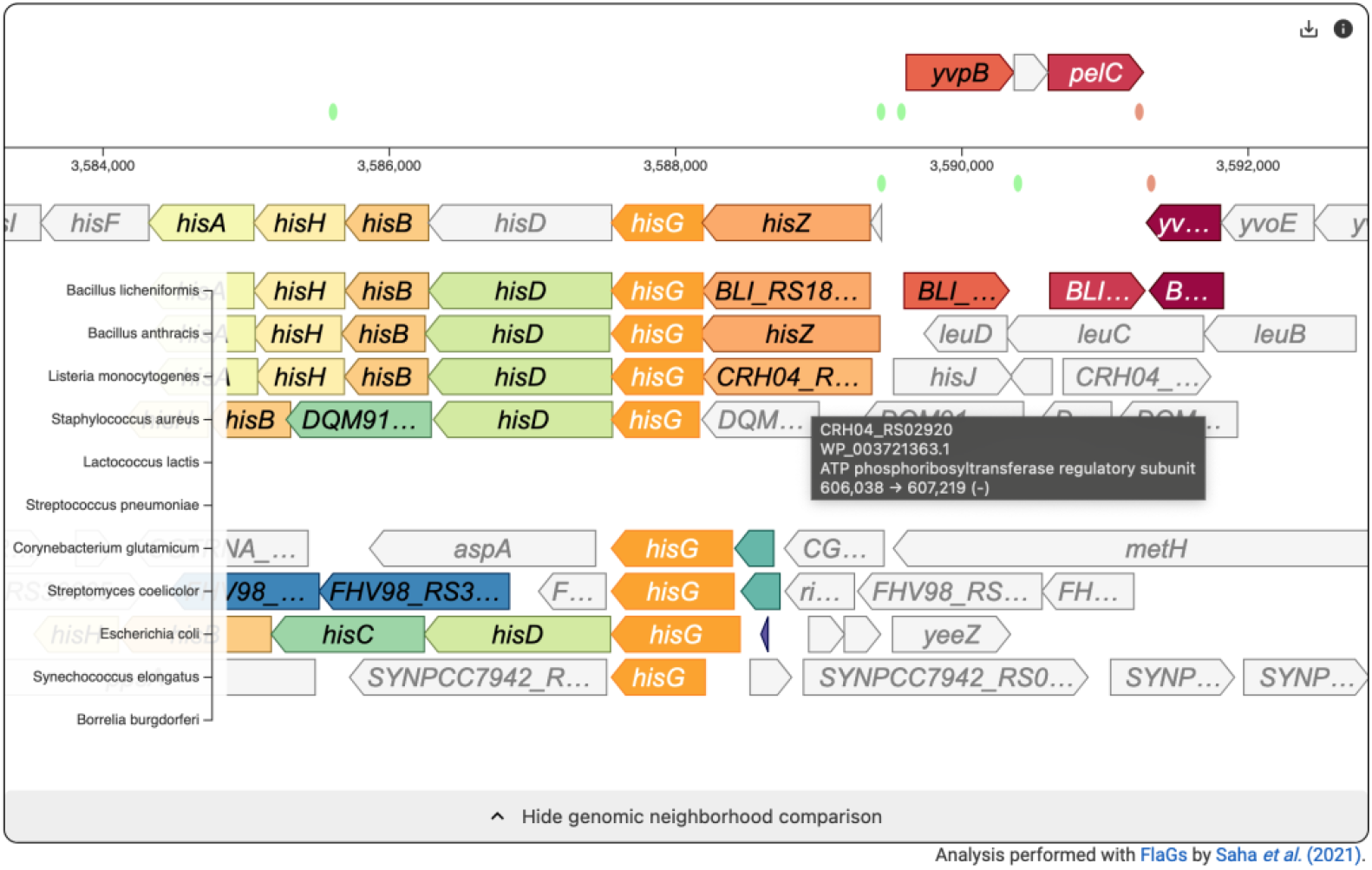
The genomic context viewer for *hisG* with expanded neighborhood comparison. The newly added neighborhood comparison allows users to explore gene order conservation across different species. Clusters of homologous genes are color-coded, and more information is displayed when hovering over a gene marker.

### New software foundation and revised API

For *Subti*Wiki’s newest version, we completely rebuilt its foundation and revised its architecture from the ground up by using established state-of-the-art frameworks. For the backend, we used Flask, a popular micro web framework written in Python. We improved the separation of data and representation by employing a REST (Representational State Transfer) interface, where the backend serves only raw data in JSON format which is visualized by the client side. For this, we used Angular, a modern frontend framework for single-page web interfaces, which is built on TypeScript and developed by Google. This new architecture allowed us to expose a public API (application programming interface), which grants direct access to *Subti*Wiki’s data for bioinformatics purposes. The REST architecture provides resources and methods to facilitate automated retrieval of information by programs and scripts. Moreover, we use Swagger to provide user-friendly documentation of the available resources and API endpoints (Fig. 5). The user interface can be accessed via the “API” link in the “Data” menu of the navigation bar.

**Figure 5.**
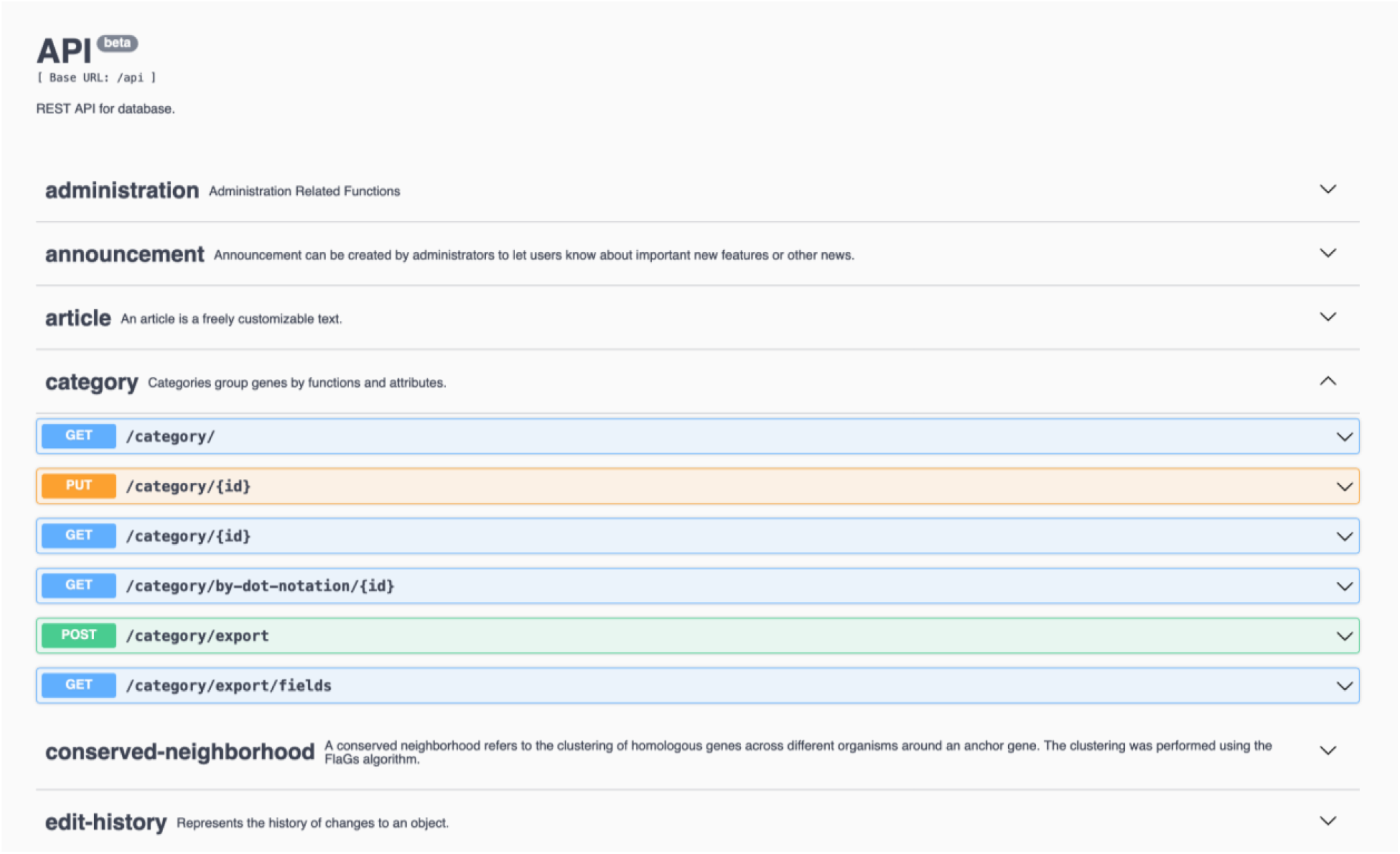
API documentation page. Resources and endpoints available via the API are documented in an interactive manner, allowing users to test different requests right in the browser.

### Future perspectives

Over the past years, *Subti*Wiki has become the most popular database that provides up-to-date and curated data for the model organism *B. subtilis*. The integration of data helps to develop novel hypotheses that can be experimentally validated, thus creating new knowledge. Given the large number of unknown proteins, this is not only very important to make *B. subtilis* one of the first organisms that are fully understood, but also for the development of biotechnological applications and to develop treatment strategies against closely related ESKAPE pathogens. Future aims include the integration of information on multiple strains of *B. subtilis* in addition to the laboratory strain 168, as well as information concerning RNA molecules as an entity on its own, with dedicated pages and information on RNA-protein and RNA-metabolite interactions. So far, *Subti*Wiki has inspired the development of additional databases for *Listeria monocytogenes, Mycoplasma pneumoniae*, and *Mycoplasma mycoides* JCVI-syn3A (24, 25, 26). With its new software foundation, *Subti*Wiki is perfectly suited as a starting point for setting up more databases for other organisms. We are confident that all this will consolidate the role of *Subti*Wiki as a trendsetter for model organism databases.

## Acknowledgements

Tom Nowak is acknowledged for his help with the development of the genomic conservation viewer. We are grateful to all lab members for testing the new features and for valuable feedback.

